# Meta-GWAS for quantitative trait loci identification in soybean

**DOI:** 10.1101/2020.10.17.343707

**Authors:** Johnathon M. Shook, Jiaoping Zhang, Sarah E. Jones, Arti Singh, Brian W. Diers, Asheesh K. Singh

## Abstract

We report a meta-Genome Wide Association Study involving 73 published studies in soybean (*Glycine max* L. [Merr.]) covering 17,556 unique accessions, with improved statistical power for robust detection of loci associated with a broad range of traits. *De novo* GWAS and meta-analysis were conducted for composition traits including fatty acid and amino acid composition traits, disease resistance traits, and agronomic traits including seed yield, plant height, stem lodging, seed weight, seed mottling, seed quality, flowering timing, and pod shattering. To examine differences in detectability and test statistical power between single- and multi-environment GWAS, comparison of meta-GWAS results to those from the constituent experiments were performed. Using meta-GWAS analysis and the analysis of individual studies, we report 483 quantitative trait loci (QTL) at 393 unique loci. Using stringent criteria to detect significant marker trait associations, 66 candidate genes were identified, including 17 candidate genes for agronomic traits, 19 for seed related traits, and 33 for disease reaction traits. This study identified potentially valuable candidate genes that affect multiple traits. The success in narrowing down the genomic region for some loci through overlapping mapping results of multiple studies is a promising avenue for community-based studies and plant breeding applications.

## INTRODUCTION

Genome-wide association studies (GWAS) analyze the association between a trait of interest and thousands of genetic variants throughout the genome. The general approach has benefited from the development of greatly increased numbers of markers due to the advent of next-generation sequencing approaches (Rico *et al*. 2013), and increased sample size with the formation of biobanks, such as the 100,000 Genomes Project (The 100,000 Genomes Project 2019). Plant scientists now routinely conduct GWAS in crop species, including soybean *[Glycine max* (L.) Merr.]. Increased marker data availability and development of new statistic methods provided great opportunities to gain new knowledge from existing data and address previous lacuna of GWAS experiments (Zeng *et al*. 2017; Chang *et al*. 2016; Chang and Hartman 2017; Bandillo *et al*. 2015; Bandillo *et al*. 2017; Zhou *et al*. 2015; de Azevedo Peixoto *et al*. 2017; Zhang *et al*. 2015; Zhang *et al*. 2017).

Researchers have recognized that while single environment GWAS such as those conducted in the greenhouse are powerful for genetic studies and candidate gene identification, their extrapolation in field environment applications require further validation (Zhang *et al*. 2015; de Azevedo Peixoto *et al*. 2017; Coser *et al*. 2017). When comparing separate studies of the same trait, significant differences in results are often found. These differences may be caused by allele frequency variation between populations, inadequate control of population structure, or environmental dependencies (Gibson and Mullen 1996). With the availability of standardized marker data across the USDA soybean germplasm collection (Song *et al*. 2015), several studies have mapped important major effect quantitative trait loci (QTL) using historical records and GWAS analysis: for example, insect resistance (Chang and Hartman 2017), disease resistance (Chang *et al*. 2016), descriptive traits such as flower and pubescence color (Bandillo *et al*. 2017), and seed oil and protein content (Bandillo *et al*. 2015). However, for many quantitative traits such as seed composition or plant height, using raw measurements from differing environments introduces bias, which may erode the power of detection for significant QTL (Chen *et al*. 2010). While results from within the same environment(s) share a common environmental component, attempting to combine multiple panels grown in different environments leads to an improper assignment of environmental effects to the differences between genetics of the panels involved (Zhao *et al*. 2019). Meta-analysis provides an attractive alternative to address the above-mentioned challenges of individual GWAS, and this analysis can be performed on results from independent studies using statistical approaches such as those provided by the analysis program METAL (Willer *et al*. 2010).

Quantitative traits, in contrast with qualitative traits, are controlled by many genes and environmental factors. To fully understand the pathways that determine these traits, interactions between previously discovered genes and new candidate genes must be added to the existing models. Directly measured traits often comprise only a portion of the information about a biological pathway, necessitating the identification of pleiotropic effects (on correlated traits) for an increased biological understanding of the phenotype. Genes may exhibit pleiotropy either through control of a common pathway such as the influence of *Dt1* on both plant height and lodging (Diers *et al*. 2018), or through multiple effects of a chemical as seen in the effect of *T* locus that has a dual role in pigmentation and chilling tolerance through isoflavones (Takahashi and Asanuma 1996). Identifying genes that control multiple phenotypes of importance can either suggest candidates for fixation, in cases where both effects are positive, or may identify possible penalties associated with incorporating particular alleles and improve multi-trait selection results (Bolormaa *et al*. 2014).

Meta-analyses include separately analyzing each individual experiment in order to determine experiment-specific p-value and allele effect estimates, rather than performing a combined analysis to leverage extensive data (Bandillo *et al*. 2015). Further genetic insights can be gleaned through an ease in the identification of pleiotropic effects due to the analysis of a wide range of traits. Moreover, the ability to compare the results from a combined analysis with those from separate analyses of individual studies allows for the identification of both environment-dependent associations and for the enrichment and detection of quantitative traits and rare alleles from more unique but diverse populations. Previous meta-analysis results have shown the effectiveness of combined panels to identify minor genes that were missed in a single study (Chang *et al*. 2017). Due to the need for adequate representation of minor alleles in GWAS, rare alleles that are predominant in a small zone of adaptation may be absent or undetectable within individual studies. The agronomic screenings for the USDA soybean germplasm collection are arranged based on the influx of new germplasm into the United States, and therefore serve as a semi-randomized subset of global soybean variation and spatiotemporal patterns in the origins of new accessions enabling potential detection of rare variants, which may be enriched in one of these geographical regions (Trotta *et al*. 2016).

While combined analyses for disease and insect resistance (Chang *et al*. 2016; Chang and Hartman 2017) and seed composition (Bandillo *et al*. 2015) have previously been reported, we perform a large-scale meta-analysis utilizing individual studies in soybean. Our study builds on previous studies by integrating the environmental component that can provide a historical perspective on adaptation, with the inclusion of quantitative traits of agronomic importance, stress tolerance, and seed composition. Subsequent study of pleiotropic genes and reporting on gene rich clusters can be useful when attempting to introgress favorable alleles into breeding lines (Cameron *et al*. 2017), as it improves the understanding of potential complications of introgression. The multitude of traits examined with our study facilitates the detection of co-localized peaks indicative of potential pleiotropic effects of genes across a diverse range of phenotypes. Loci associated with multiple traits identified within this study require additional functional validation, as GWAS are not designed to definitively differentiate between pleiotropy and (tight) linkage. We included results from reports published from 1964 to 2009 for a total of 73 individual studies. The design of this study was intended to identify co-localization of peaks for multiple traits, as well as to identify previously overlooked genes through meta-analysis approaches. Using meta-GWAS analysis and analysis of individual studies, we report 393 unique QTL including 66 candidate genes across important traits and provide confirmation of many previously reported genes. This study provides targets for functional characterization and introgression of previously untapped diversity for many important traits.

## MATERIALS AND METHODS

### Genotypic data and quality control

Marker data from the testing of 20,087 *Glycine max* and *G*. *soja* accessions from the USDA Soybean Germplasm Collection with the SoySNP50K iSelect BeadChip (Song *et al*. 2013) were downloaded from SoyBase (Song *et al*. 2013). A data imputation pipeline based on Java implementation of Beagle 5.0 (Browning and Browning 2016) was utilized to impute missing data for the 42,080 SNP markers that were aligned to the Williams 82 reference genome v2 assembly. Markers aligned to scaffolds but not assigned to a chromosome were removed prior to processing. Ten burn-in iterations and five phasing iterations were used to impute missing markers, which accounted for 0.64% of all markers. For each test, markers remaining after applying cutoffs of minor allele frequency ≥ 0.05 for studies involving 300 ≤n ≤1000 accessions, or 0.01 for studies involving n ≥ 1001 accessions, were selected for further analysis.

### Phenotypic data and genetic accessions

Numeric phenotypic data from USDA reports were compiled from the U.S. National Plant Germplasm System website (http://npgsweb.ars-grin.gov/gringlobal/descriptors.aspx (Descriptors for Soybean 2019). Subsets of accessions that were part of historical USDA germplasm characterization trials with a size n ≥ 300 were selected for further analysis. Information on the design of the original trials is available from the technical bulletins in which they were originally published. These technical bulletins are available online in part at https://pubs.nal.usda.gov/sites/pubs.nal.usda.gov/files/tb.htm (Miller 2003). Alternatively, PDFs of the technical bulletins are available on our GitHub (https://github.com/SoylabSingh/META-GWAS). Additional traits, such as disease resistance and amino acid composition, were downloaded from the NPGS website.

### Genome-wide association analysis

Each experiment was analyzed separately with a mixed linear model implemented using GAPIT in R (Lipka *et al*. 2012) to prevent confounding of environmental effects with marker effects, which would be expected for several traits (i.e., flowering time, oil, protein, etc.). Population structure was controlled using the first three PCAs based on the marker data. This resulted in 585 combinations of experiment/trait analyses. Analysis was subsequently performed for combined panels for each trait. The Bonferroni threshold (Neyman and Pearson 1928) was employed to minimize the likelihood of false positives in declaring significance. The significant SNPs were compiled for further analysis (**Supplemental Table 1**).

Initial QTL calling was performed trait-by-trait based on marker position. Subsequently, QTL for related traits (such as flowering date and maturity date) with substantial overlap were merged, resulting in fewer unique QTL than originally called. Local LD decay analysis was used to further clarify between separate or overlapping QTL.

Markers that were significant for multiple traits and experiments, or were identified during analysis of the combined trials, were examined for nearby candidate genes. Candidate genes were identified by examining annotated genes within linkage disequilibrium (LD) of the leading SNP with r^2^ > 0.7 for each experiment and peak (de Azevedo Peixoto *et al*. 2017). Candidate gene identification was performed based on previously characterized genes, gene family function, and the nearest gene to the peak SNP in cases where no known function could be identified. For candidate casual genetic variant analysis, we utilized the SNP dataset from the genome resequencing study of 302 soybean lines (Zhou *et al*. 2015) and searched the possible causal mutants at the identified candidate genes. We first identified the lead SNP from peaks of interest in the resequencing dataset, then calculated the pairwise LD r^2^ values between the lead SNP and the SNPs covering the locus of candidate gene. All other analyses here within were aligned to the Glyma2.0 reference genome (https://soybase.org/gb2/gbrowse/gmax2.0/). The R package ‘circlize’ was employed to generate the circular visualizations of significant SNPs for multiple traits throughout the genome. Study names have been shortened for convenience within the text; a reference file is provided to find the initial source of phenotypic data used in this work (**Supplemental Table 2**). Trait definitions, as well as the number of QTL and candidate genes identified for each trait, are provided in **Supplemental Table 3**.

## RESULTS AND DISCUSSION

From the individual study GWAS and meta-GWAS 4,919 significant SNPs were detected, of which 787 were reported from the meta-GWAS analysis. Complete listing of the significant SNP identified using individual study GWAS and meta-GWAS are provided in **Supplemental Table 1**. Among these 787 SNPs identified using meta-GWAS, 110 were associated with agronomic traits, 106 with seed composition traits, and 571 with disease resistance traits. Overall, candidate genes were assigned for 66 unique loci; and these included genes with moderate to large effects. We focus our results on loci that were associated with multiple traits.

### Agronomic Traits

Amongst agronomic traits, we identified 1422 marker-trait associations with traditional GWAS studies, as well as 110 SNPs associated with agronomic traits when analyzed across studies by meta-GWAS. In all, 115 QTL across 20 chromosomes were identified, with 17 candidate genes (**Figure 1a, Supplemental Table 1, Supplemental Table 3, Table 1**).

**Figure 1.**
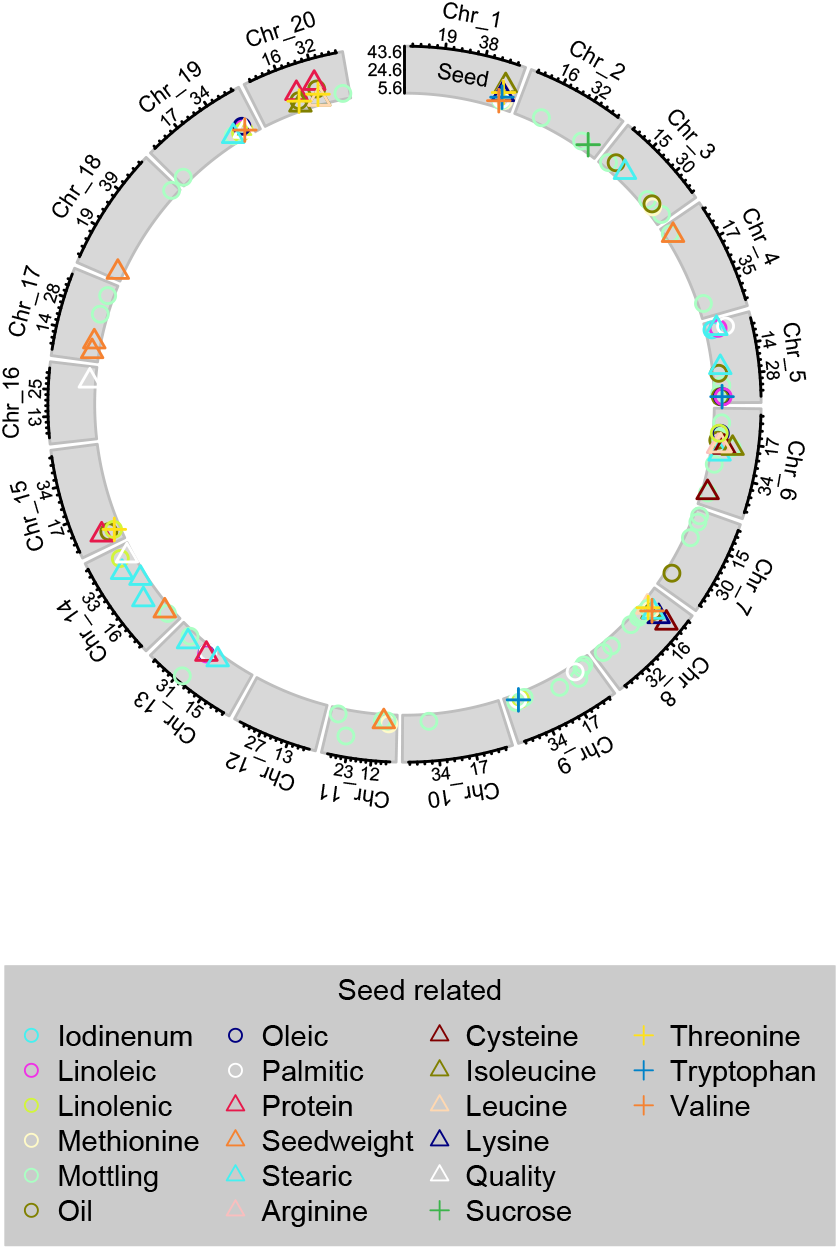
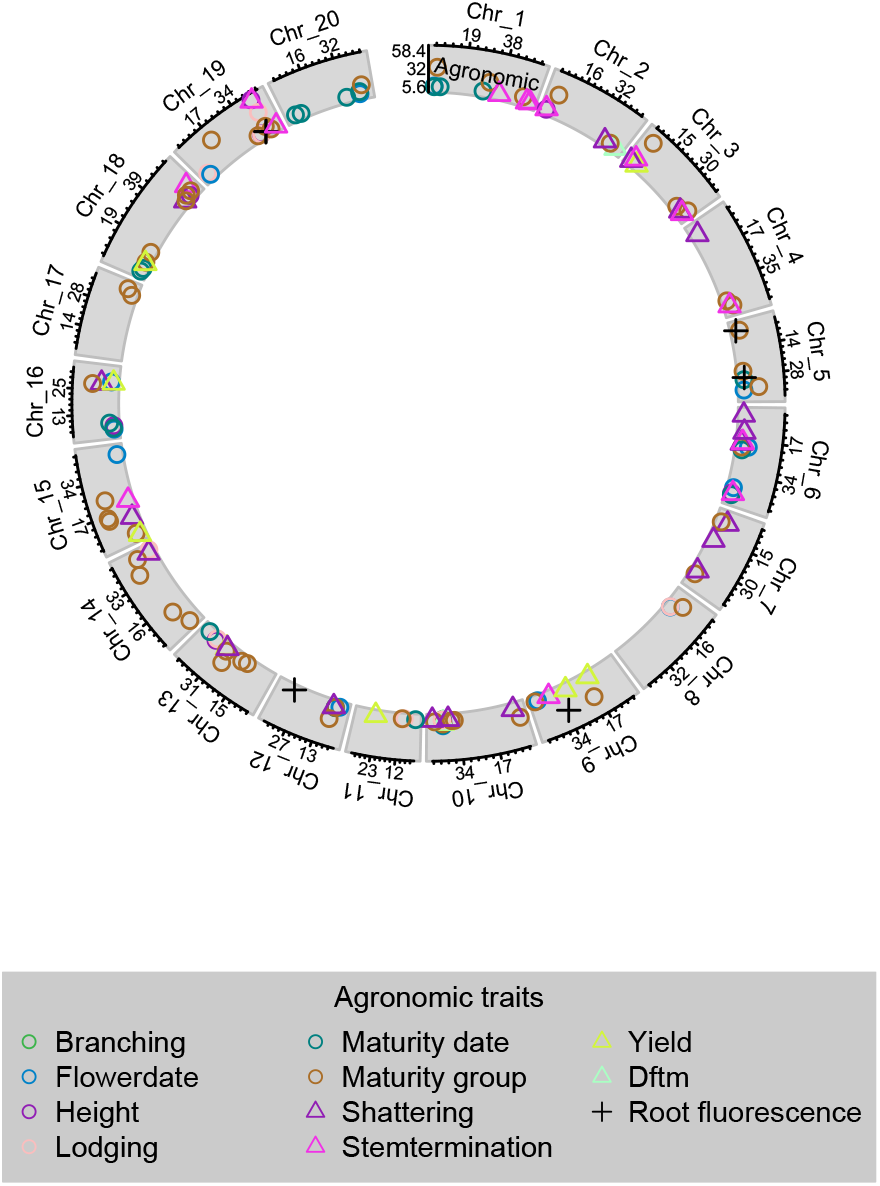
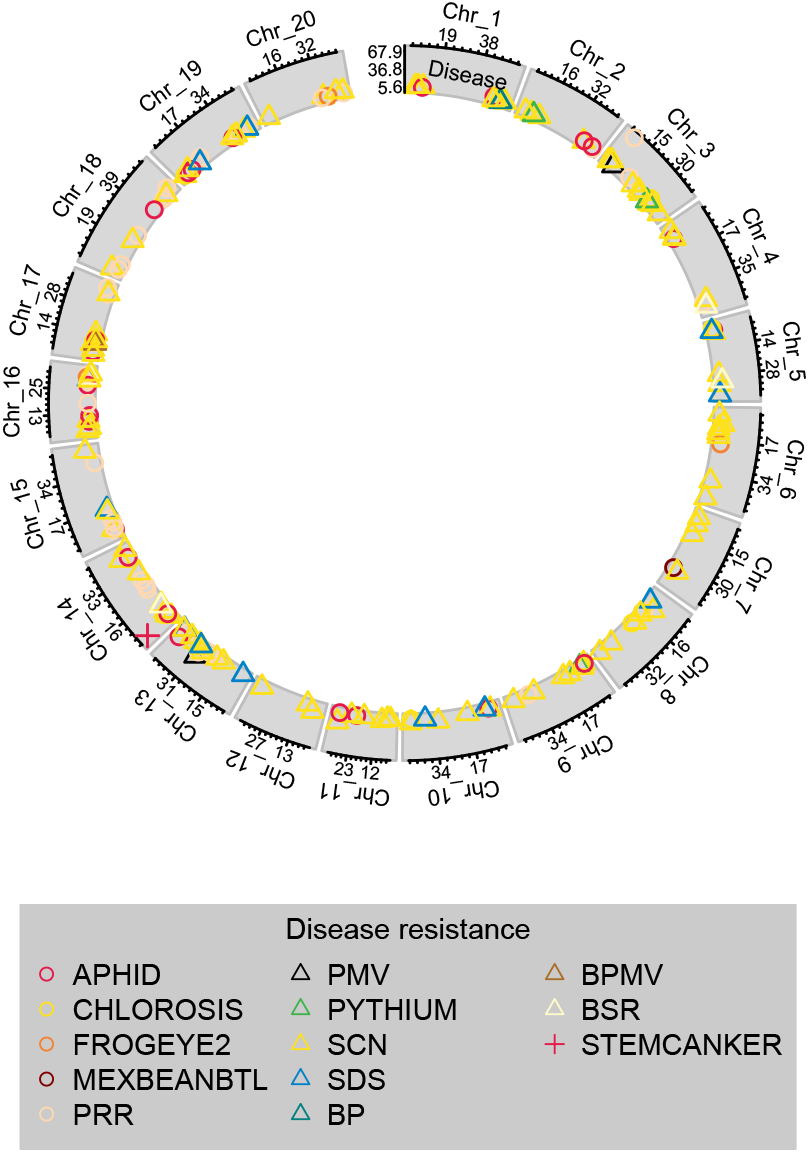
Significant SNPs from GWAS from individual studies and meta-GWAS. (a) Peaks for seed related traits, (b) Peaks for flowering and maturity related traits, (c) Peaks for disease resistance related traits. Symbol position along the x-axis shows the position (in Mb) along the chromosome, while y-axis symbol position shows the LOD score of the lead SNP for each QTL. X-axis labels indicate position (in Mb) of tertile points, while y-axis labels show minimum, maximum, and middle of LOD score range for the given trait class. Shape and color correspond to unique traits.

**Table 1.** List of candidate genes identified for agronomic traits using GWAS from individual studies and Meta-GWAS.

| Chromosome | Likely Gene | Meta-GWAS | Individual Studies GWAS | Trait(s) | Studies Source |
| --- | --- | --- | --- | --- | --- |
| 5 | *Glyma.05G200100* |  | * | Flower date, Maturity date, Maturity group | 4il87, ms1999.01, ms923 |
| 6 | *E1* | * | * | Flower date, Maturity date, Maturity group, Stem termination | 1il64, 1il66, 2il81.1, 2il81.2, 4il87, 5il90, il0102, il989, meta, mn945 |
|  | *Glyma.06G068900* | * | * | Seed mottling | 3mn83.2, meta |
|  | *Glyma.06g134400* |  | * | Pod shattering (early), Pod shattering (late) | 4il87 |
|  | *T* | * | * | Seed mottling | 3il84, meta, ms1999.01, ms923, ms967 |
| 7 | *Glyma.07g049800* | * | * | Pod shattering (early), Pod shattering (late) | 3il84, meta, ms1999.01, ms923 |
| 8 | *I* |  | * | Seed mottling | 1il66, 2ky81, 4il87, il0102, ms923 |
| 9 | *fr1* |  | * | Root fluorescence | fluorjt97 |
|  | *Glyma.09g090600* | * | * | Seed mottling | 1il66, 4il87, meta |
|  | *Glyma.09g266200* |  | * | Flower date, Maturity group | ms923, ms1999.01 |
| 10 | *E2* | * | * | Branching, Flower date, Height, Maturity date, Maturity group, Yield | 1il64, 1il66, 2il81.1, 3il83.1, 3il84, il0102, il989, meta, ms1999.01, ms967 |
| 11 | *K1/AGO* | * | * | Seed mottling | 3mn83.2, il0102, meta, ms923, ms967 |
| 13 | *Rsv1* | * | * | Seed mottling | 1il66, 2il81.1, 2il81.2, 5il90, meta, ms1999.01, ms2000.02, ms923 |
| 14 | *fan1* |  | * | Seed quality | 2ky81 |
| 15 | *Glyma.15g139800* | * | * | Pod shattering (early), Pod shattering (late) | 1il66, 2il81.2, 2ky81, meta |
| 16 | *E9* | * | * | Flower date, Maturity group | 2il81.1, 3il83.1, meta, ms1999.01 |
|  | *Pdh1* | * | * | Pod shattering (early), Pod shattering (late) | 1il64, 2il81.1, 4il87, il0102, meta, ms1999.01, ms2000.02, ms923, ms967 |
| 18 | *Dt2* | * | * | Stem termination | meta, mn945, ms923 |
| 19 | *ABC,*<br>*Glyma.*<br>*19g016*<br>*400* | * | * | Lodging | 1il66, 2ky81, ms923, 3il84, meta |
|  | *Dt1,*<br>*Glyma.*<br>*19g194*<br>*300* | * | * | Height, Lodging,<br>Stem termination | 1il64, 1il66, 2il81.1, 2il81.2, 2ky81,<br>3il83.1, 3il84, 3mn83.2, 4il87, 5il90,<br>il0102, meta, mn945, ms1999.01,<br>ms2000.02, ms923, ms967 |
|  | *E3* |  | * | Maturity group | 2il81.2 |

In our approach, we used results from individual studies to detect overlapping genomic regions for the purpose of locating candidate gene for traits, including for genes previously cloned. The locus harboring *Dt1 (Glyma.19g194300)* (Liu *et al*. 2010), the major gene conditioning stem termination in soybean, was significantly associated with oleic acid and linoleic acid content, as well as plant height, stem termination, and stem lodging (**Supplemental Table 1**). By comparing the mapping results of four studies, we were able to limit the candidate genomic region to a 125 kb fragment harboring previously cloned *Dt1* (from *ss715635422* to *ss715635460)* (**Supplemental Figure 1**). These results highlight the advantages of meta-GWAS for finer mapping the candidate gene region. A nonsynonymous SNP (*SNP19_44980087),* in high LD (r^2^ = 0.5) with the leading SNP *ss715635424* (also known as *SNP_19_45000827),* was found at the fourth exon of *Dt1* that changes amino acid R (Arg) to W (Trp) (**Supplemental Figure 2**). This SNP is identical to the R166W mutation previously identified (Liu *et al*. 2010).

On chromosome 19, we identified a QTL for stem lodging which was on the opposite end of the chromosome as *Dt1*., Stem lodging is associated with plant height and this has been reported in multiple crops (Flint-Garcia *et al*. 2003; Diers *et al*. 2018; Singh *et al*. 2019). As lodging causes significant yield and quality losses, the development of the shorter statured wheat and rice were promoted which could better handle high input agriculture. However, this solution is not universally applicable. In soybean, pods are arranged at nodes on the stem and decreasing the length of stem, and if fewer nodes are present, yield potential is reduced. Leveraging four studies, we report a peak for tolerance to stem lodging with the candidate gene *Glyma.19g016400*, an ABC transporter on chromosome 19. This locus was found to affect lodging tolerance but was not found to be associated with plant height, thereby making it a useful target to develop lodging resistant soybean cultivars without decreasing stem length and yield potential. While this is the first genome wide association study identifying this gene, additional evidence towards its validity comes from several recent patents (US Patents #8697941, 8748695, and 9675071) that relate to molecular markers in the region of interest and include *Glyma.19g016400* as one of the candidate genes for PPO inhibitor tolerance in soybean. Significant effects of this region for seed yield, lodging, and plant height were reported from the SoyNAM project (Diers *et al*. 2018). The results from Hulting *et al*. (2001) on PPO inhibitor tolerance and our findings on stem lodging susceptibility suggests a tradeoff between PPO inhibitor tolerance and lodging susceptibility. The soybean accessions highly tolerant to sulfentrazone contain alleles associated with increased lodging in our study, necessitating further studies to validate these observations.

On chromosome 6, a significant SNP peak was identified that co-located with the *T* gene, a flavonoid 3’ hydroxylase (Toda *et al*. 2002). This region was significant for arginine, cysteine, isoleucine, and leucine levels, as well as for seed mottling (**Figure 1 a, c**). The cloned *E2* locus (Watanabe *et al*. 2011) was significantly associated with flowering and maturity date, maturity group, days from flowering to maturity, plant height, and seed yield (**Figure 1b**). The associations between *E2* and these traits has been previously reported (Fang *et al*. 2017).

### Seed Composition Traits

Amongst seed composition traits, we identified 1364 marker-trait associations with traditional GWAS studies, as well as 106 SNPs associated with compositional traits when analyzed across studies by meta-GWAS. SNPs associated with composition were found on chromosomes 1-9, 11, 13-15, 17, 19-20, resulting in 88 QTL with 19 candidate genes (**Figure 1b, Supplemental Table 1, Supplemental Table 3, Table 2**)

A cluster of candidate genes for seed composition, including isoleucine, methionine, leucine, tryptophan, threonine, lysine, and palmitic acid, were located in a region of 30 kb on chromosome 1 between 53.13 – 53.16 Mb, 4 a cysteine desulfurase (*Glyma.01g197100)* and a malate and lactate dehydrogenase gene (*Glyma.01g197700)* (**Supplemental Figure 1**). Further targeted analysis will be necessary to determine which gene is influencing each trait, as a single enzyme is unlikely responsible for multiple steps in the metabolic pathway. We found significant SNPs in high LD (r^2^ > 0.5) with the detected leading SNP at the promoter of *Glyma.01g197700,* but not in the coding region of the gene (**Supplemental Figure 2**).

A region including the *I* locus on chromosome 8 (Clough *et al*. 2004) was associated with seed mottling, as well as oil, cysteine, isoleucine, leucine, linoleic acid, lysine, methionine, palmitic acid, stearic acid, threonine, and valine levels in the seed (**Figure 1b**). The most likely candidate gene for the observed differences in amino acids levels, *AK-HDSH* (aspartokinase homoserine dehydrogenase, *Glyma.08g107800)* is a bifunctional enzyme catalyzing the key steps of asparagine phosphatization and the aspartate-semialdehyde to homoserine conversion by which aspartate family amino acids (lysine, threonine, methionine, and isoleucine) are synthesized (Zhu-Shimoni and Galili 1998). However, amino acid data were generated using Near Infrared Reflectance, which may have low precision in estimating amino acid composition when there is variability in seed coat color (Baianu *et al*. 2011). Therefore, further validation is needed to establish the association between the *AK-HDSH* or *I* loci and the amino acid profile.

*SACPD-C (Glyma.14g121400)* was the primary candidate to explain differences in stearic acid content within seed oil and has been previously functionally validated (Gillman *et al*. 2014). Using the Wm82.a2 reference genome build, this appeared as three separate peaks; however, a single peak was observed when using the Wm82.a1 version. We postulate a possible assembly error in the region surrounding the *SACPD-C* locus in the soybean reference genome Wm82.a2, due to conflicting results (**Supplemental Table 4**). We attempted to identify false peaks generated due to genome mis-assembly by fitting the lead SNP as a covariate in the GWAS model, and then observed lower p-values for the remaining SNPs and detected a weaker signal from surrounding SNPs indicative of a single gene. Presence of stronger signals in surrounding SNPs would have indicated that two separate genes are in play. Additionally, the r^2^ between SNPs in all three regions was greater than 0.7, suggesting physical linkage. The Wm82.a1 results (SNP effects, physical location, LD) provide the most plausible explanation for the presence of a single gene in this genomic region and suggests that Wm82.a2 has unresolved errors in scaffold positioning.

A peak on chromosome 5 associated with palmitic acid content was detected in 3 different studies. Using data from the ‘2mn81’ study, the locus mapped to a region of over 600 kb. However, two other studies (2ky81 and ms2000.02) mapped this locus within a smaller region of 130 kb (*ss715592495-ss715592503)* and 182 kb (*ss715592491-ss715592500),* respectively, with an overlap of about 88 kb (*ss715592495*-*ss715592500*) (**Supplemental Figure 2**). The candidate gene *FATB1a* (*Glyma.05g012300*) (Wilson *et al*. 2001) was identified in the overlap. However, no SNP in LD (r^2^ ≥ 0.5) with the leading SNP of the locus was identified at the coding region or promoter of *FATB1a* based on analysis of resequencing data (Zhou et al. 2015) except the synonymous *SNP_5_7995427* (**Supplemental Figure 1**). Causal variants have been identified in mutagenized breeding material (Thapa *et al*. 2016, Bachleda *et al*. 2016. Goettel *et al*. 2016), but naturally occurring variations are not well characterized.

### Disease Resistance Traits

Amongst disease traits, we identified 1346 marker-trait associations with traditional GWAS studies, as well as 571 SNPs associated with disease traits when analyzed across studies by meta-GWAS. 213 QTL mapped to all 20 chromosomes, with 33 candidate genes identified (**Figure 1c**, **Supplemental Table 1, Supplemental Table 3, Table 3**). Meta-analysis in several instances narrowed the genomic region for QTL. For example, the association between the *Rps3* region and resistance to race 1 of *Phytophthora* root rot was mapped to a 144 kb region in the meta-analysis, compared to a 1Mb region in individual studies (**Supplemental Table 1**). This reduces the search space for causal genes and allows for greater accuracy when identifying candidate genes.

**Table 2.**
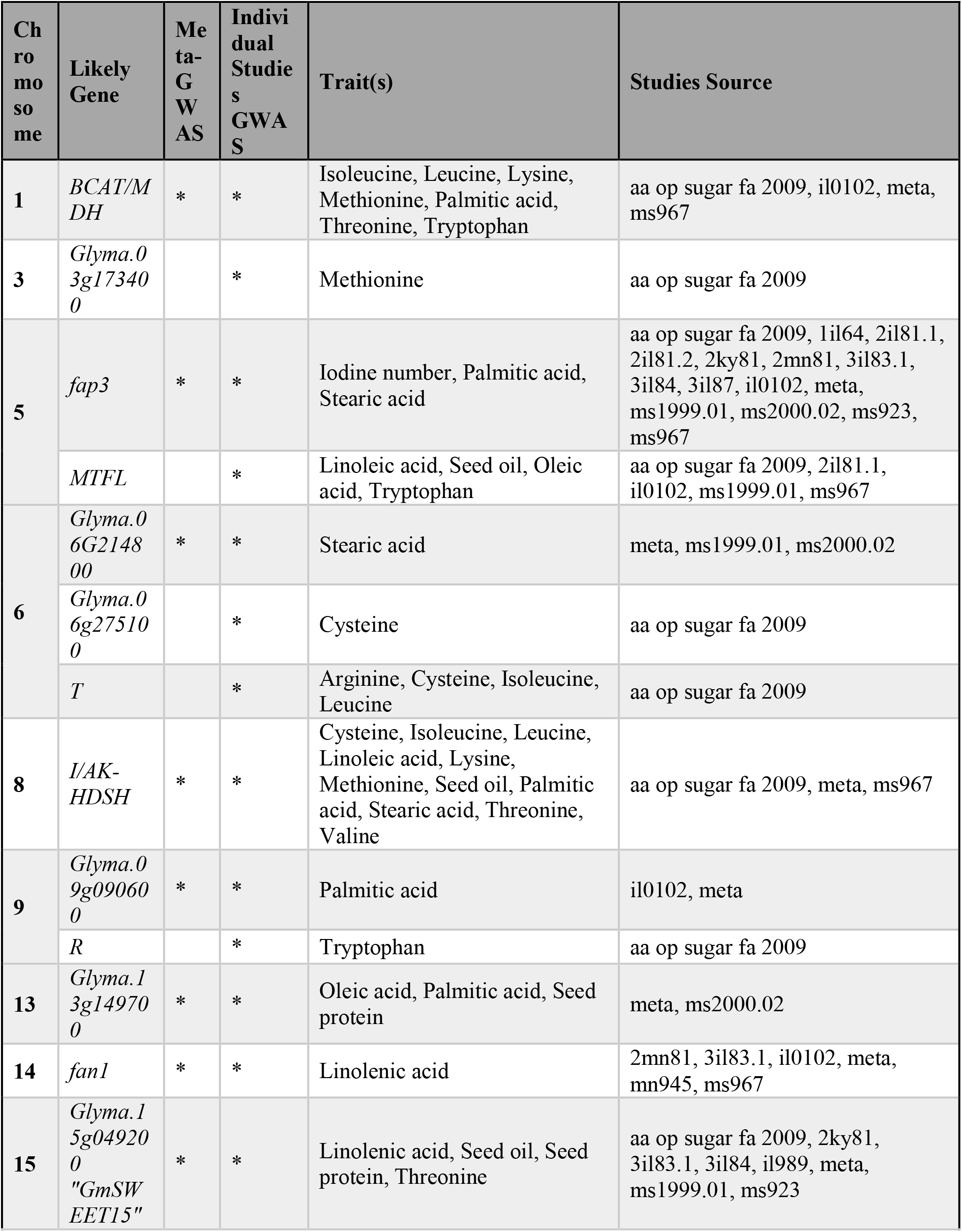
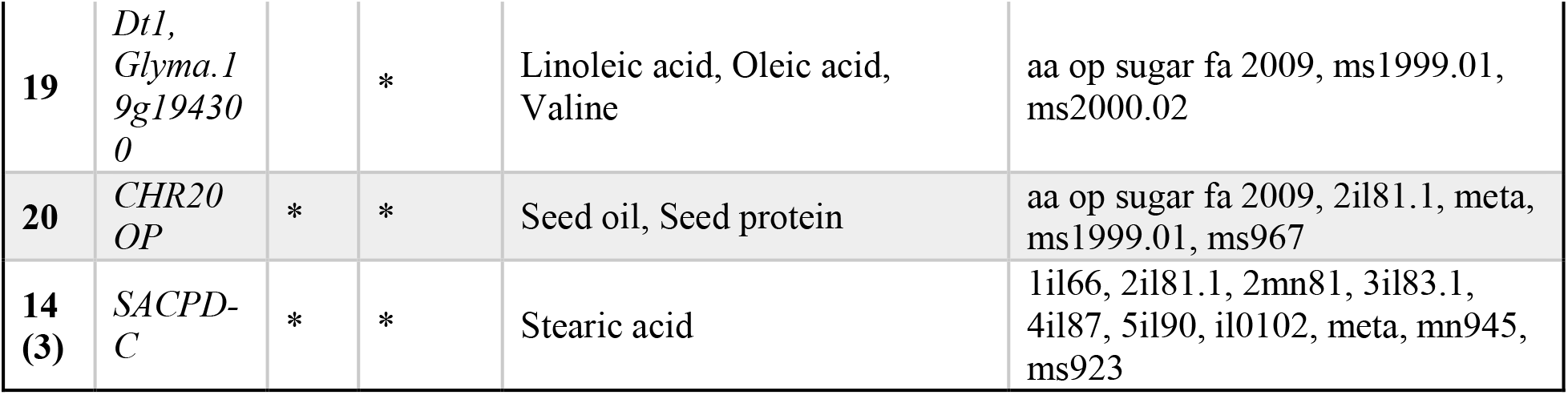
List of candidate genes identified for seed composition traits using GWAS from individual studies and Meta-GWAS.

**Table 3.**
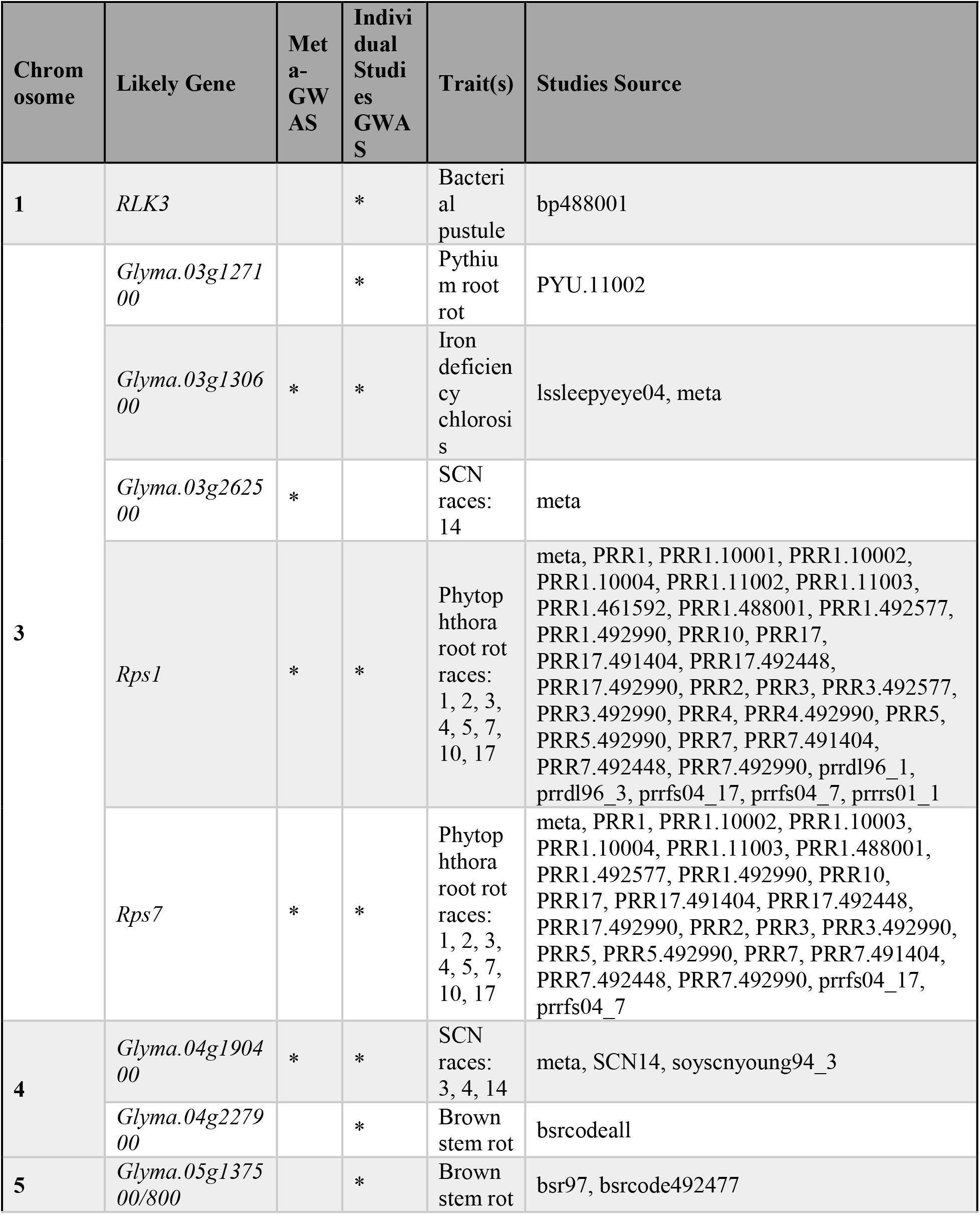
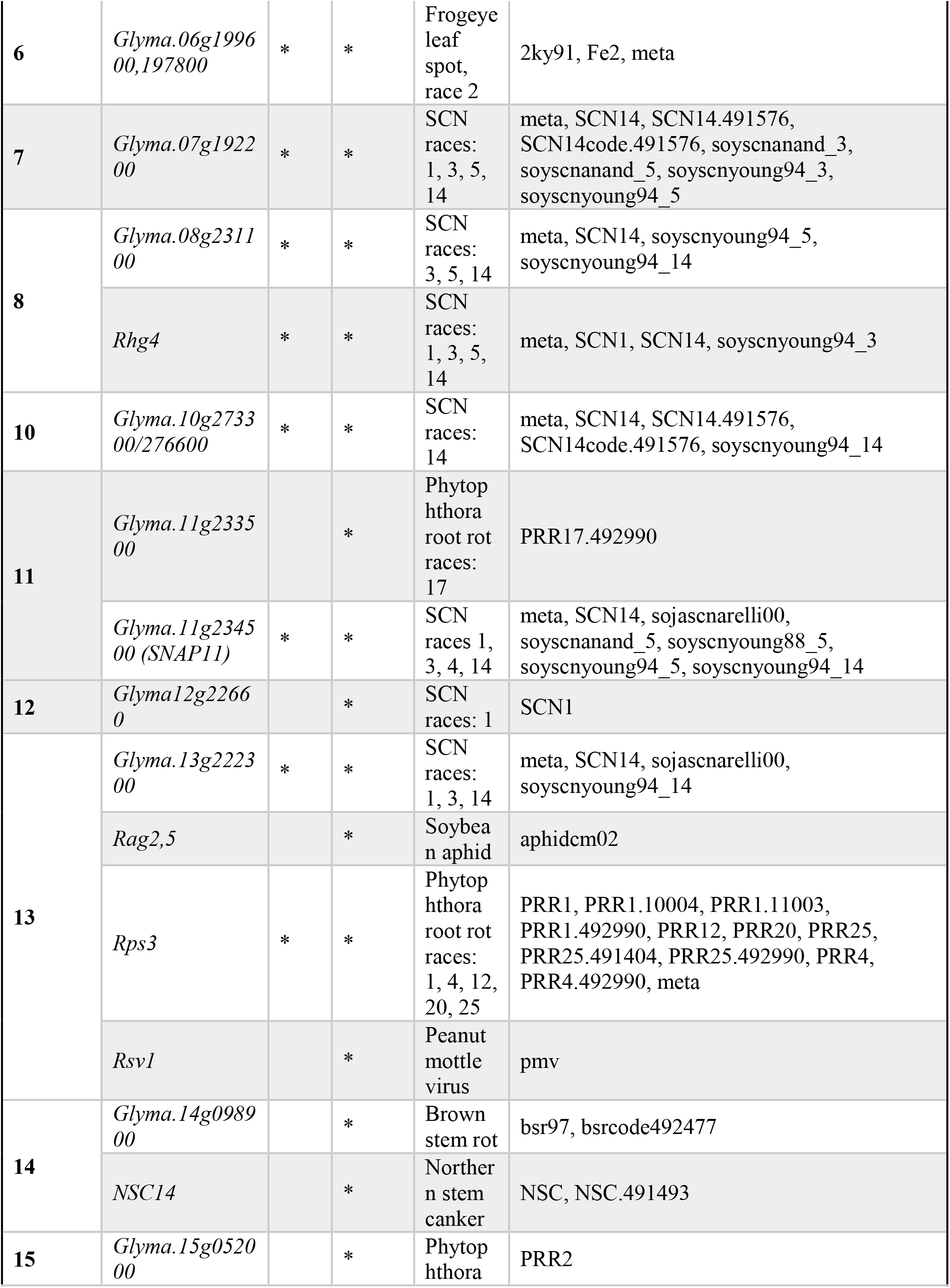
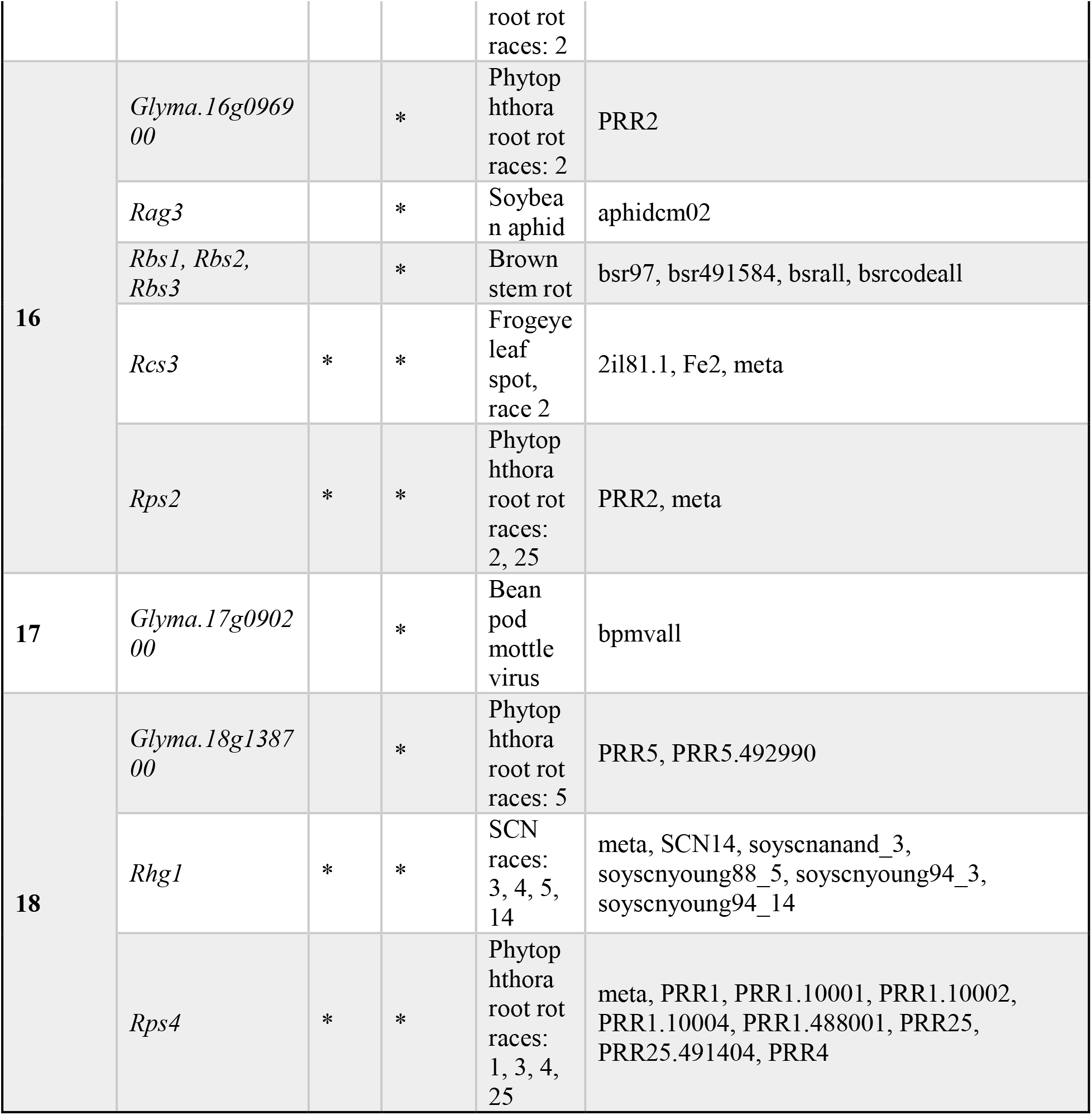
List of candidate genes identified for disease resistance/ stress tolerance traits using GWAS from individual studies and Meta-GWAS.

We found a peak that was associated with resistance to races 1, 2, 3, 4, 5, 7, 10, and 17 of *Phytophthora sojae* that mapped to the position of the *Rps1* locus (Gao and Bhattacharyya 2008). A previously unreported peak for soybean cyst nematode resistance was identified on chromosome 11 was mapped to *Glyma.11g234500,* an alpha-soluble N-ethylmaleimide-sensitive factor (NSF) attachment protein (α-SNAP). Notably, the candidate genes *GmSNAP11 (Glyma.11g234500)* and *GmSNAP14 (Glyma.14g054900)* (Lakhssassi et al. 2017), identified at 7 kb and 84 kb apart from lead SNPs *ss715610420* and *ss715618859*, respectively, are paralogs and encode a Soluble NSF Attachment Protein (SNAP). Another soybean SNAP gene on chromosome 18, *GmSNAP18*, has been reported to play a role in resistance to SCN (Cook et al. 2012). On chromosome 1, the locus for seed composition co-localized with a bacterial pustule resistance QTL. This QTL does not correspond to the previously identified *Rxp* locus, instead, a candidate gene Glyma.01g197800 is identified as the potential underlying gene. A peak on chromosome 3 at 34.24 −35.18 Mb was found to be significantly associated with iron deficiency chlorosis tolerance and *Pythium irregulare* resistance. This region has previously been investigated as the source of IDC tolerance in “Isoclark” (Stec et al. 2013). The GWAS analysis identified previously unreported genomic regions that were associated with resistance to bean pod mottle virus, brown stem rot, frogeye leaf spot, Phytophthora root rot, and soybean cyst nematode (**Figure 1c**). A full list of identified SNPs and candidate genes for these traits, as well as for all other traits examined in this study using both combined analyses and analysis of individual experiments are provided in **Supplemental Table 1**.

The majority of studies included in this work for disease resistance were germplasm screenings, where many entries were tested to find new sources of resistance. Such germplasm screening studies were not originally intended for GWAS; for example, multiple rating systems, ordinal rating scales, and noninteger ratings used in the studies complicates result comparisons and are not easily amenable to linear statistical models. Standardization of screening protocols across research groups and inclusion of key data for comparison of studies such as those suggested by the MIAPPE checklist (Ćwiek-Kupczyńska et al. 2016) will be key for future research into plant disease resistance. In addition, an increased utilization of image-based phenotyping will play a key role, allowing for digital disease severity ratings on a continuous scale (Naik et al. 2017; Zhang et al. 2017), minimal inter- and intra-rater variability in measurements through hyperspectral camera and ML-based analysis (Nagasubramanian et al. 2018; Nagasubramanian et al. 2019). It will also enable the comparison of results across studies by facilitating reanalysis of previous experiments with new rating systems or approaches, as long as needed input variables are available.

### Implications of pleiotropy vs. linked genes

While repeated crossing or careful selection of the donor parent can break linkage drag, negative pleiotropic effects associated with a gene of interest are more problematic. Candidate gene analysis was aided by tissue-specific gene expression data available at SoyBase. The use of a blend of individual and meta-analyses provided improved resolution through examining overlapping peaks and utilizing the increased power in larger panels in the meta-analysis. When investigating the peak on chromosome 1 for fatty acid and amino acid composition, a convincing distinction between pleiotropy and linkage could not be made. This was due to the presence of multiple strong candidate genes. While meta-GWAS approaches are very beneficial for improving map resolution, they are still limited in their inference in regions with strong linkage disequilibrium. Meta-GWAS results outputs still require follow-up molecular and functional validation to confirm the candidate genes as well as to confirm pleiotropy vs. linkage.

Pleiotropic effects of major genes significantly alter multiple traits simultaneously, creating a situation of either rapid improvement across traits, or of tradeoffs, such as is found in most soybean protein/oil content QTL. Genetic improvement utilizing pleiotropic effects may be limited in applicability to specific geographic regions if they affect key adaptation genes such as the maturity loci or stem termination. Therefore, it will be necessary for breeders to independently determine whether a gene with pleiotropic effects is a good fit for their variety development goals. In cases where pleiotropy is associated with a tradeoff between multiple traits, such as between seed protein and oil content, breeders will need to weigh the importance of each trait or identify combinations of genes affecting the trait that can provide an adequate phenotype for each trait considered.

### Motivation for the use of meta-analysis

For many important row crop species, such as soybean, corn, wheat, and sorghum, it is impractical or impossible to evaluate the full breadth of the available germplasm at a single location. This is due to space limitations, availability of labor or funding for phenotyping, or irreconcilable differences between genotypes preventing them from growing in the same place, such as differences in photoperiod sensitivity or vernalization requirements. To capture the breadth of the genetic and phenotypic diversity, it is necessary to test each variety with a similarly adapted cohort. The separate analysis of each environment can increase the odds of finding alleles which are near fixation in the population or are environmentally dependent (Singh et al. 2014; Sherman et al. 2019).

For simple, qualitative traits such as pubescence color in soybean, there is little benefit in meta-GWAS due to the consistency with which the gene can be mapped and the lack of environmental dependence on trait expression. When studying environmentally dependent traits, such as agronomic, disease resistance and seed composition traits including seed oil or protein content, meta-GWAS provide advantages particularly in increasing the likelihood of finding small effect genes. When comparing individual experiments results (**Figure 2a**) with the combined meta-analysis (**Figure 2b**), additional significant peaks were observed in meta-analysis. For example, the SNP marker *ss715614263* was previously associated with seed protein using mega-analysis (Bandillo et al. 2015). The same locus was found to be associated with protein, palmitic, and oleic acid content in an individual panel in the current study (ms2000.02), but was associated with protein and linoleic acid content in the meta-analysis (**Supplemental Table 1**). While meta-analysis identified fewer traits in the specific instance of *ss715614263,* the association with an additional trait (compared to individual analysis) still encourages its use, as each newly associated trait may provide guidance in identifying putative causal genes. A full listing of candidate genes detected in each study is provided as **Supplemental Table 5,** which also provides a reference to candidate genes detected either only in individual studies or only via meta-analysis. Identification of an association with multiple related traits, although only spanning one to two markers, is a strong signal that the association may merit additional study to identify a strong candidate gene and further explore the possible pleiotropic effects this locus is exhibiting, especially when a stringent cut-offs are used to declare significance.

**Figure 2.**
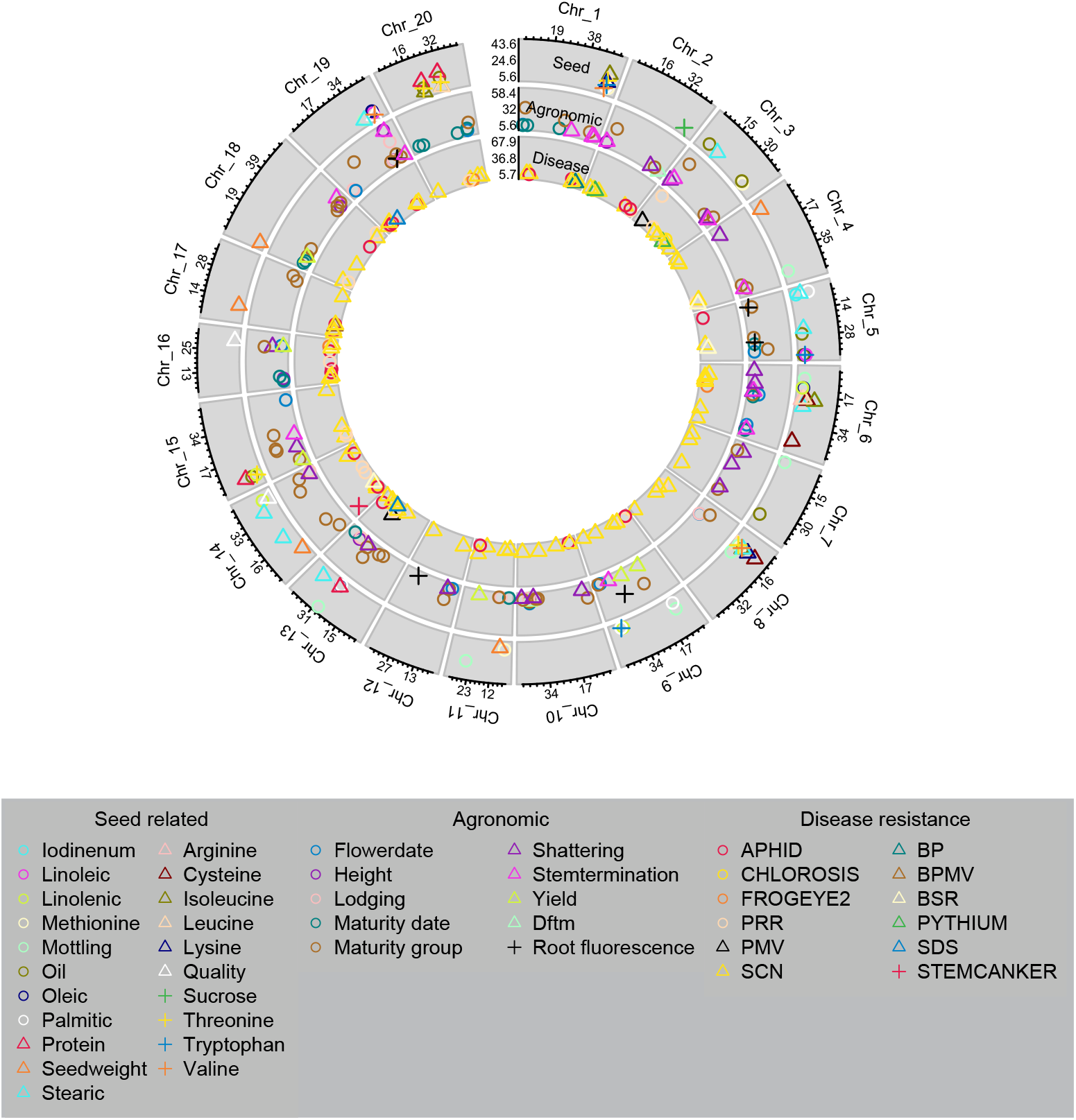
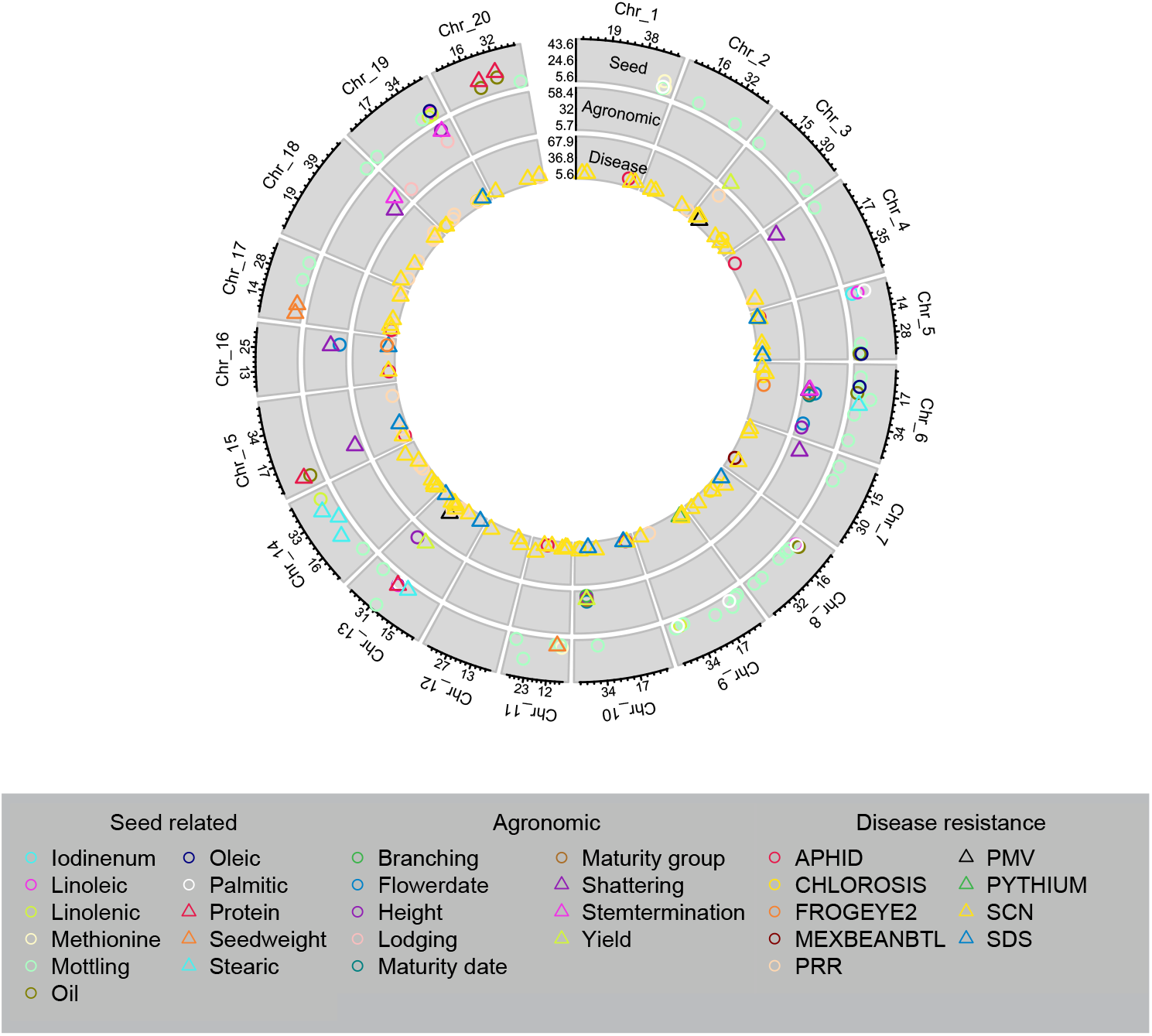
Circle plots of significant SNPs identified with (a) GWAS from individual studies, and (b) meta-GWAS. The peaks in the innermost ring includes seed composition traits, the middle ring includes disease resistance traits, and the outermost ring includes agronomic traits. Symbol position along the x-axis shows the position (in Mb) along the chromosome, while y-axis symbol position shows the LOD score of the lead SNP for each QTL. X-axis labels indicate position (in Mb) of tertile points, while y-axis labels show minimum, maximum, and middle of LOD score range for the given trait class. Shape and color correspond to unique traits

To maximize the effectiveness of soybean breeding programs, we sought to identify as many genes as possible for numerous traits, ensuring that multiple paths are available for further cultivar improvement. By maximizing the identified links between markers and phenotypes of interest, meta-GWAS aids efforts to bridge the gap between genotype and phenotype, allowing for improvements not only in trait prediction and selections, but also in modelling the interactions between multiple genes in overall trait performance.

### Future mapping, validation and integration with Phenomics studies

Traditional fine mapping through creating lines sharing homogenous genetic background, such as near isogenic lines, is a powerful tool to uncover the casual genetic variants. However, it is time consuming to develop new near-isogenic lines in multiple backgrounds to reduce the potential influence of background-specific effects. In this study, large variation of LD architecture was observed across populations. This enables substantially shortening of the candidate chromosomal regions of specific QTL by comparing mapping results from separate studies using different populations. Considering almost all accessions in the USDA Soybean Germplasm Collection were genotyped by SoySNP50K BeadChip and are publicly accessible, mapping populations with a high LD decay rate at specific genomic regions of interest can be constructed for fine mapping. The consistent identification of major genes, including those affecting multiple traits of interest, suggests that further improvements in mapping ability would likely require a model with the major genes treated as covariates. While it is currently possible to account for the effects of major genes by using SNPs linked to the gene of interest as covariates, this approach is only an approximation due to incomplete linkasge between common SNPs and the underlying gene. Instead, allelespecific markers should be developed and deployed across both wild-type germplasm and breeding material.

In the future, similar studies will benefit by incorporating weather, soil, or management parameters in order to explain differences in marker effects between individual studies and in Meta-GWAS (Cook et al. 2017). In this scenario, access to standardized, quality-controlled records will be needed to tease apart the GxE component and identify the architecture of environmentally mediated expression and decipher associations between genetics and environmental signals for the traits of interest. The establishment of standardized tests enabled with advanced sensors and high-throughput phenotyping should improve the opportunity to identify additional genes influencing traits of interest through the analysis of previously ignored component traits, such as leaf expansion rate or chlorophyll density in the case of yield, (Dhondt et al. 2013) which may lead to an increased understanding of the genetic architecture of these traits and responses to environmental and management conditions (Parmley et al. 2019).

## CONCLUSION

Combined analysis of all investigated traits found 63 loci that corresponded to previously reported QTL, characterized genes, and new reported loci backed up with strong candidate genes conditioning the observed phenotypes. Several of the previously identified loci (for example, *Dt1, E2*) were associated with multiple traits, identifying putative pleiotropic effects of the underlying genes. Differences between results in individual trials and the combined analyses confirm the importance of multi-environment testing for identification of key traits, but also provide a strong motivation to create a community database that can be queried for scientific advancement. Continued publication of raw phenotypic values from screenings will increase the power for identification of important genes for both mean and plastic responses to reduce the financial and time burden on any individual program while benefitting future breeders and researchers. For example, the sharing of phenotypic information across research programs both nationally and globally, as currently on-going with multi-states and –institutions uniform soybean tests and other cooperatively run tests in other crops.

## Data availability statement

The authors affirm that all data necessary for confirming the conclusions of the article are present within the article, figures, and tables. Raw data and codes will be available at https://github.com/SoylabSingh/Soy-Meta-GWAS.

## Author contribution

AS, AKS conceptualized the study; JS, AS, AKS designed the study; JS conducted statistical analysis with contributions from AKS and JZ; Figures were prepared by JZ with inputs from JS; JS interpreted the results with contributions from JZ, SJ, AS, BD, AKS; JS wrote the first draft with AKS; All authors contributed in writing, reviewing, and approve the manuscript.

## Acknowledgements

Authors sincerely thank all researchers past and present who generated data for individual studies and set up a community resource for advancing soybean research and development. We thank David Blystone and Dr. David Grant, which greatly helped the manuscript. We thank the Iowa Soybean Association (to AKS), R F Baker Center for Plant Breeding (to AKS), Monsanto Chair in Soybean Breeding (to AKS), USDA IOW04314, and National Research Traineeship (to JMS) for the financial support.

## SUPPLEMENTAL FILES

**Supplemental Figure 1.** Comparison of the chromosomal region of (a) *FATB1a,* (b) *Dt1*, (c) *PMDH1* loci identified using diverse populations. The x-axis indicates the physical location on each chromosome referring soybean genome version Glyma2.0. The y-axis indicates the pairwise LD r^2^ between the lead SNP and the rest SNPs in the specific region for each population.

**Supplemental Figure 2.** SNP at the region of candidate genes (a) *FATB1a*, (b) *Dt1*, (c) *PMDH1*. SNP were retrieved from Figshare database (http://figshare.com/articles/Soybean_resequencing_project/1176133) based on the genome resequencing study of the 302 diverse soybean lines. For each panel, the x-axis indicates the physical location of the specific regions on the chromosome. The y-axis indicates the pairwise LD r^2^ between the SNP(s) in the region and the lead SNP, which was also identified in the resequencing dataset.

**Supplemental Table 1.** Full list of significant marker-trait associations found in individual GWAS and meta-GWAS.

**Supplemental Table 2.** List of studies, methods, and reference literature used to generate phenotypic datasets.

**Supplemental Table 3.** Significant SNPs for stearic acid levels from 3il83.1. Positions in Wms82.1 and Wms82.2 provided to show alignment differences between the two reference genome versions.

**Supplemental Table 4.** Trait definitions, number of QTL detected, and number of candidate genes assigned for each trait.

**Supplemental Table 5.** Full listing of which candidate genes were detected in which study, as well as whether the association was detected in only individual studies or only in meta-analysis.

## References

Assefa, T., J. Zhang, R.V. Chowda-Reddy, A.N. Moran Lauter, A. Singh et al., 2020 Deconstructing the genetic architecture of iron deficiency chlorosis in soybean using genome-wide approaches. BMC Plant Biology 20 (1):42.

Bachleda, N., Pham, A. & Li, Z. Identifying *FATB1α* deletion that causes reduced palmitic acid content in soybean N87-2122-4 to develop a functional marker for marker-assisted selection. Mol Breeding 36, 45 (2016). https://doi.org/10.1007/s11032-016-0468-9

Baianu, I., J. Guo, R. Nelson, T. You, and D. Costescu, 2011 NIR Calibrations for Soybean Seeds and Soy Food Composition Analysis: Total Carbohydrates, Oil, Proteins and Water Contents [v.2] Nat Preceedings. https://doi.org/10.1038/npre.2011.6611.1

Bandillo, N., D. Jarquin, Q. Song, R. Nelson, P. Cregan et al., 2015 A Population Structure and GenomeWide Association Analysis on the USDA Soybean Germplasm Collection. The Plant Genome 8 (3).

Bandillo, N.B., A.J. Lorenz, G.L. Graef, D. Jarquin, D.L. Hyten et al., 2017 Genome-wide Association Mapping of Qualitatively Inherited Traits in a Germplasm Collection. The Plant Genome 10 (2).

Bernard, R.L., 1972 Two genes affecting stem termination in soybeans. Crop Science 12:235–239.

Bolormaa, S., J.E. Pryce, A. Reverter, Y. Zhang, W. Barendse et al., 2014 A Multi-Trait, Meta-analysis for Detecting Pleiotropic Polymorphisms for Stature, Fatness and Reproduction in Beef Cattle. PLOS Genetics 10 (3):e1004198.

Browning, Brian L., and Sharon R. Browning, 2016 Genotype Imputation with Millions of Reference Samples. Am J Hum Genet 98 (1):116–126.

Cameron, J.N., Y. Han, L. Wang, and W.D. Beavis, 2017 Systematic design for trait introgression projects. Theor Appl Genet 130 (10):1993–2004.

Chang, D., M.A. Nalls, I.B. Hallgrímsdóttir, J. Hunkapiller, M. van der Brug et al., 2017 A meta-analysis of genome-wide association studies identifies 17 new Parkinson’s disease risk loci. Nat Genet 49 (10):1511–1516.

Chang, H.-X., and G.L. Hartman, 2017 Characterization of Insect Resistance Loci in the USDA Soybean Germplasm Collection Using Genome-Wide Association Studies. Frontiers in Plant Science 8 (670).

Chang, H.-X., A.E. Lipka, L.L. Domier, and G.L. Hartman, 2016 Characterization of Disease Resistance Loci in the USDA Soybean Germplasm Collection Using Genome-Wide Association Studies. Phytopathology™ 106 (10):1139–1151.

Chen, X., F. Zhao, and S. Xu, 2010 Mapping environment-specific quantitative trait loci. Genetics 186 (3):1053–1066.

Clough, S.J., J.H. Tuteja, M. Li, L.F. Marek, R.C. Shoemaker et al., 2004 Features of a 103-kb gene-rich region in soybean include an inverted perfect repeat cluster of CHS genes comprising the I locus. Genome 47 (5):819–831.

Cook, D.E., T.G. Lee, X. Guo, S. Melito, K. Wang et al., 2012 Copy number variation of multiple genes at Rhgl mediates nematode resistance in soybean. Science 338 (6111):1206–1209.

Cook, J., Mahajan, A. & Morris, A. Guidance for the utility of linear models in meta-analysis of genetic association studies of binary phenotypes. Eur J Hum Genet 25, 240–245 (2017). https://doi.org/10.1038/ejhg.2016.150

Coser, S.M., R.V. Chowda Reddy, J. Zhang, D.S. Mueller, A. Mengistu et al., 2017 Genetic Architecture of Charcoal Rot (Macrophomina phaseolina) Resistance in Soybean Revealed Using a Diverse Panel. Frontiers in Plant Science 8 (1626).

de Azevedo Peixoto, L., T.C. Moellers, J. Zhang, A.J. Lorenz, L.L. Bhering et al., 2017 Leveraging genomic prediction to scan germplasm collection for crop improvement. PLOS ONE 12 (6):e0179191.

Descriptors for Soybean, 2019. U.S. National Plant Germplasm System.

Dhondt, S., N. Wuyts, and D. Inzé, 2013 Cell to whole-plant phenotyping: the best is yet to come. Trends Plant Sci 18 (8):428–439.

Diers, B.W., J. Specht, K.M. Rainey, P. Cregan, Q. Song et al., 2018 Genetic Architecture of Soybean Yield and Agronomic Traits. G3: Genes| Genomes|Genetics 8 (10):3367–3375.

Fang, C., Y. Ma, S. Wu, Z. Liu, Z. Wang et al., 2017 Genome-wide association studies dissect the genetic networks underlying agronomical traits in soybean. Genome Biol 18 (1):161.

Flint-Garcia, S.A., C. Jampatong, L.L. Darrah, and M.D. McMullen, 2003 Quantitative Trait Locus Analysis of Stalk Strength in Four Maize Populations Mention of a trademark or proprietary product does not constitute a guarantee, warranty, or recommendation of the product by the USDA or the University of Missouri, and does not imply its approval to the exclusion of others that may be more suitable. Crop Science 43 (1):13–22.

Gao, H., and M.K. Bhattacharyya, 2008 The soybean-Phytophthora resistance locus Rps1-k encompasses coiled coil-nucleotide binding-leucine rich repeat-like genes and repetitive sequences. BMC Plant Biol 8:29.

Gibson, L.R., and R.E. Mullen, 1996 Soybean seed composition under high day and night growth temperatures. Int J Mol Sci 73 (6):733–737.

Gillman, J.D., M.G. Stacey, Y. Cui, H.R. Berg, and G. Stacey, 2014 Deletions of the SACPD-C locus elevate seed stearic acid levels but also result in fatty acid and morphological alterations in nitrogen fixing nodules. BMC Plant Biol 14 (1):143.

Goettel W, Ramirez M, Upchurch RG, An YQ. Identification and characterization of large DNA deletions affecting oil quality traits in soybean seeds through transcriptome sequencing analysis. Theor Appl Genet. 2016;129(8):1577–1593. doi:10.1007/s00122-016-2725-z

Gu, Z., L. Gu, R. Eils, M. Schlesner, and B. Brors, 2014 circlize implements and enhances circular visualization in R. Bioinformatics 30 (19):2811–2812.

Hulting, A.G., L.M. Wax, R.L. Nelson, and F.W. Simmons, 2001 Soybean (Glycine max (L.) Merr.) cultivar tolerance to sulfentrazone. Crop Protection 20 (8):679–683.

Lakhssassi, N., S. Liu, S. Bekal, Z. Zhou, V. Colantonio et al., 2017 Characterization of the Soluble NSF Attachment Protein gene family identifies two members involved in additive resistance to a plant pathogen. Sci Rep 7 (1):45226.

Lipka, A.E., F. Tian, Q. Wang, J. Peiffer, M. Li et al., 2012 GAPIT: genome association and prediction integrated tool. Bioinformatics 28 (18):2397–2399.

Liu, B., S. Watanabe, T. Uchiyama, F. Kong, A. Kanazawa et al., 2010 The Soybean Stem Growth Habit Gene *Dt1* Is an Ortholog of Arabidopsis TERMINAL FLOWER1. Plant Physiology 153 (1):198.

Miller, E.K., 2003 Index to USDA Technical Bulletins, edited by USDA/ARS. National Agricultural Library.

Nagasubramanian, K., S. Jones, S. Sarkar, A.K. Singh, A. Singh et al., 2018 Hyperspectral band selection using genetic algorithm and support vector machines for early identification of charcoal rot disease in soybean stems. Plant Methods 14 (1):86.

Nagasubramanian, K., S. Jones, A.K. Singh, S. Sarkar, A. Singh et al., 2019 Plant disease identification using explainable 3D deep learning on hyperspectral images. Plant Methods 15 (1):98.

Naik, H.S., J. Zhang, A. Lofquist, T. Assefa, S. Sarkar et al., 2017 A real-time phenotyping framework using machine learning for plant stress severity rating in soybean. Plant Methods 13 (1):23.

Neyman, J., and E.S. Pearson, 1928 On the use and interpretation of certain test criteria for purposes of statistical inference, Part I. Biometrika 20A (1-2):175–240.

Parmley, K., K. Nagasubramanian, S. Sarkar, B. Ganapathysubramanian, and A.K. Singh, 2019 Development of Optimized Phenomic Predictors for Efficient Plant Breeding Decisions Using Phenomic-Assisted Selection in Soybean. Plant Phenomics 2019:15.

Sherman, R.M., J. Forman, V. Antonescu, D. Puiu, M. Daya et al., 2019 Assembly of a pan-genome from deep sequencing of 910 humans of African descent. Nat Genet 51 (1):30–35.

Singh, A., R.E. Knox, R.M. DePauw, A.K. Singh, R.D. Cuthbert et al., 2014 Stripe rust and leaf rust resistance QTL mapping, epistatic interactions, and co-localization with stem rust resistance loci in spring wheat evaluated over three continents. Theor Appl Genet 127 (11):2465–2477.

Singh, D., X. Wang, U. Kumar, L. Gao, M. Noor et al., 2019 High-Throughput Phenotyping Enabled Genetic Dissection of Crop Lodging in Wheat. Frontiers in Plant Science 10 (394).

Song, Q., D.L. Hyten, G. Jia, C.V. Quigley, E.W. Fickus et al., 2013 Development and Evaluation of SoySNP50K, a High-Density Genotyping Array for Soybean. PLOS ONE 8 (1):e54985.

Song, Q., D.L. Hyten, G. Jia, C.V. Quigley, E.W. Fickus et al., 2015 Fingerprinting Soybean Germplasm and Its Utility in Genomic Research. G3: Genes| Genomes| Genetics 5 (10):1999.

Srour, A., A.J. Afzal, L. Blahut-Beatty, N. Hemmati, D.H. Simmonds et al., 2012 The receptor like kinase at Rhg1-a/Rfs2 caused pleiotropic resistance to sudden death syndrome and soybean cyst nematode as a transgene by altering signaling responses. BMC genomics 13:368–368.

Stec, A.O., P.B. Bhaskar, Y.-T. Bolon, R. Nolan, R.C. Shoemaker et al., 2013 Genomic heterogeneity and structural variation in soybean near isogenic lines. Frontiers in plant science 4:104–104.

Takahashi, R., and S. Asanuma, 1996 Association of T gene with chilling tolerance in soybean. Crop Science 36:559–562.

Thapa, R., Carrero-Colón, M. and Hudson, K.A. (2016), New Alleles of *FATB1A* to Reduce Palmitic Acid Levels in Soybean. Crop Science, 56: 1076–1080. doi:10.2135/cropsci2015.09.0597

The 100,000 Genomes Project, 2019. GenomicsEngland.

Toda, K., D. Yang, N. Yamanaka, S. Watanabe, K. Harada et al., 2002 A single-base deletion in soybean flavonoid 3’-hydroxylase gene is associated with gray pubescence color. Plant Mol Biol 50 (2):187–196.

Trotta, L., T. Hautala, S. Hämäläinen, J. Syrjänen, H. Viskari et al., 2016 Enrichment of rare variants in population isolates: single AICDA mutation responsible for hyper-IgM syndrome type 2 in Finland. Eur J Hum Genet 24 (10):1473–1478.

Watanabe, S., Z. Xia, R. Hideshima, Y. Tsubokura, S. Sato et al., 2011 A Map-Based Cloning Strategy Employing a Residual Heterozygous Line Reveals that the GIGANTEA Gene Is Involved in Soybean Maturity and Flowering. Genetics 188 (2):395.

Willer, C.J., Y. Li, and G.R. Abecasis, 2010 METAL: fast and efficient meta-analysis of genomewide association scans. Bioinformatics 26 (17):2190–2191.

Zeng, A., P. Chen, K. Korth, F. Hancock, A. Pereira et al., 2017 Genome-wide association study (GWAS) of salt tolerance in worldwide soybean germplasm lines. Mol Breeding 37 (3):30.

Zhang, J., H.S. Naik, T. Assefa, S. Sarkar, R.V.C. Reddy et al., 2017 Computer vision and machine learning for robust phenotyping in genome-wide studies. Sci Rep 7 (1):44048.

Zhang, J., A. Singh, D.S. Mueller, and A.K. Singh, 2015 Genome-wide association and epistasis studies unravel the genetic architecture of sudden death syndrome resistance in soybean. The Plant Journal 84 (6):1124–1136.

Zhang, J., and A.K. Singh, 2020 Genetic Control and Geo-Climate Adaptation of Pod Dehiscence Provide Novel Insights into Soybean Domestication. G3: Genes|Genomes|Genetics 10 (2):545.

Zhao, J., C. Sauvage, J. Zhao, F. Bitton, G. Bauchet et al., 2019 Meta-analysis of genome-wide association studies provides insights into genetic control of tomato flavor. Nat Comm 10 (1):1534.

Zhou, Z., Y. Jiang, Z. Wang, Z. Gou, J. Lyu et al., 2015 Resequencing 302 wild and cultivated accessions identifies genes related to domestication and improvement in soybean. Nat Biotech 33 (4):408–414.

Zhu-Shimoni, J.X., and G. Galili, 1998 Expression of an Arabidopsis Aspartate Kinase/Homoserine Dehydrogenase Gene Is Metabolically Regulated by Photosynthesis-Related Signals but Not by Nitrogenous Compounds. Plant Physiology 116 (3):1023–1028.

Ćwiek-Kupczyńska, H., T. Altmann, D. Arend, E. Arnaud, D. Chen et al., 2016 Measures for interoperability of phenotypic data: minimum information requirements and formatting. Plant Methods 12 (1):44.

